# STN-PFC circuit related to attentional fluctuations during non-movement decision-making

**DOI:** 10.1101/2023.12.10.571030

**Authors:** Shengnan Ai

## Abstract

Decision-making is a cognitive process, in which participants need to attend to relevant information and ignore the irrelevant information. Previous studies have described a set of cortical areas important for attention. It is unclear whether subcortical areas also serve a role. The subthalamic nucleus (STN), a part of basal ganglia, is traditionally considered a critical node in the cortico-basal ganglia-thalamus-cortico network. Given the location of the STN and its widespread connections with cortical and subcortical brain regions, the STN plays an important role in motor and non-motor cognitive processing. We would like to know if STN is also related to fluctuations in attentional task performance, and how the STN interacts with prefrontal cortical regions during the process. We examined neural activities within STN covaried with lapses of attention (defined as behavior error). We found that decreased neural activities in STN were associated with sustained attention. By examining connectivity across STN and various sub-regions of the prefrontal cortex (PFC), we found that decreased connectivity across areas was associated with sustained attention. Our results indicated that decreased STN activities were associated with sustained attention, and the STN-PFC circuit supported this process.

**Significance Statement:** Attention is a core internal state variable that governs the allocation of limited resources depending on the task demands in various cognitive processes. If the subcortical area, subthalamic nucleus (STN), related to attentional fluctuations, and how the STN interacted with cortical regions during the process remains unclear. In this study, we examined neural activities within STN, and connectivity between STN and various prefrontal sub-regions during sustained attention and lapses of attention. We found both neural activities within STN and connectivity between STN-PFC circuit decreased during sustained attention. These findings indicated that decreased STN activities were associated with sustained attention, and the STN-PFC circuit supported this process.

## Introduction

Attention is a core internal state variable that governs the allocation of limited resources depending on the task demands in various cognitive processes. Building on decades of functional neuroimaging research, two widespread cortical networks were characterized, including the dorsal attention network (DAN) and the salience network (SN) (Dubey, et al., 2023; Corbetta et al., 2002). The DAN is believed as neural basis of top-down attention: goals at higher levels inform the focus of attention at lower levels, while SN is believed as neural basis of bottom-up attention: stimulus at lower levels informs decisions regarding which information to attend to (Suzuki et al., 2013; Gregoriou, et al., 2014; Buckley et al., 2009; Buschman et al., 2015; Foxe et al., 2011; Seeley et al., 2007; Uddin et al., 2014). These two networks interact dynamically with each other during attentional processing (Vossel et al., 2005).

Besides advances in neuroimaging technologies, the ability to conduct high-density multi-channel electroencephalogram (EEG) and magnetoencephalography (MEG) have unprecedentedly progressed our understanding of the physiology of human selective attention. Findings from neurophysiological studies indicated that theta and BHG activities held important roles in attentional process. EEG and MEG oscillation study in human revealed that theta power in dorsolateral, ventral, and medial frontal regions is related to attentional selection during cognitive conflict processing (Haciahmet et al., 2021; McDermott et al., 2017). Electrocorticography (ECoG) study found increases in high gamma (HG) power (70-250 Hz) compared to baseline over frontal, parietal, and visual areas. The high gamma power increases were modulated by the phase of the delta/theta (2-5 Hz) oscillation during attentional allocation (Szczepanski et al., 2014). iEEG study also found that theta power (∼4 Hz) mediated fluctuations in widespread cortical areas are behaviorally related to visual attention performance, and high-frequency band (HFB; 70-150 Hz) activity was nested in an ongoing 4 Hz oscillation (Helfrich et al., 2016).

These attention-dependent changes in neurophysiology have so far mainly focused on cortical regions in the brain. It is unclear whether subcortical areas also serve a role. The STN is a small lens-shaped subcortical nucleus as a part of basal ganglia. It is traditionally considered a critical node in the cortico-basal ganglia-thalamus-cortico network, which composed of the direct pathway, indirect pathway, and hyper-direct pathway (Drummond et al., 2020; Nambu et al., 2002; Eliasmith et al., 2012). Given the location of the STN and its widespread connections with cortical and subcortical brain regions (Taylor et al., 2007; Coxon et al., 2012; Forstmann et al., 2012; King et al., 2011), the STN plays an important role in motor and non-motor cognitive processing (Zavala et al., 2018; Wessel et al., 2019; Herz et al., 2017; Haynes et al., 2013; Popov et al., 2013; Kühn et al., 2005). We would like to know if STN is also related to fluctuations in attentional task performance, and how the STN interacts with cortical regions during the process.

In the present study, we address these unanswered questions by examining neural activities within STN and connectivity across regions located in three prefrontal regions and STN covaried with lapses of attention (defined as behavior errors), as has been suggested in previous studies (Weissman et al., 2006; Eichele et al., 2008; Kucyi et al., 2020). Our results showed that theta/beta power activities, ITPC, and theta-BHG PAC in STN tended to decrease during sustained attention. Besides, decreased PLV and theta-BHG PAC across areas were found during sustained attention. By further examining the spectral Granger causality across STN and prefrontal regions, we found significant STN leading information flow to prefrontal regions in the theta band, which may be the driving force of the activity in prefrontal regions during the task. Our results indicated that decreased STN activities were associated with sustained attention, and the STN-PFC circuit supported this process by decreasing the connectivity between areas.

## Materials and methods

The dataset we used was from research (Zavala et al., 2017). The information of participants, task, iEEG recordings, and electrode localization was reported in detail in (Zavala et al., 2017). Here, we introduce them in brief.

### Participants

18 patients undergoing deep brain stimulation (DBS) surgery for Parkinson’s disease participated. This study used data from all participants for behavioral analysis, and data from 9 participants for local field potential analysis, in order to get relatively balanced correct and incorrect trials.

### Behavioral task

Participants performed a novel working memory task with non-motor decision-making during the task (Fig. 1A). Each task session involved 30 blocks. At the beginning of each block, an image of square or octagon was displayed on the screen to indicate the target shape for the upcoming block. During each block, participants were sequentially presented with eight randomly chosen single-digit numbers within a square or an octagon, and instructed to attend to and remember the numbers within the target shape. Each number was displayed for 500 ms, following a blank screen for 500 ± 100 ms. During the presentation of the numbers, participants were instructed to limit all movements if possible. At the end of each block, the participants were prompted to retrieve the four target numbers vocally. Fourteen participants performed one session while LFP recordings were captured from the left STN and a second session while LFP recordings were captured from the right STN.

**Fig. 1.**
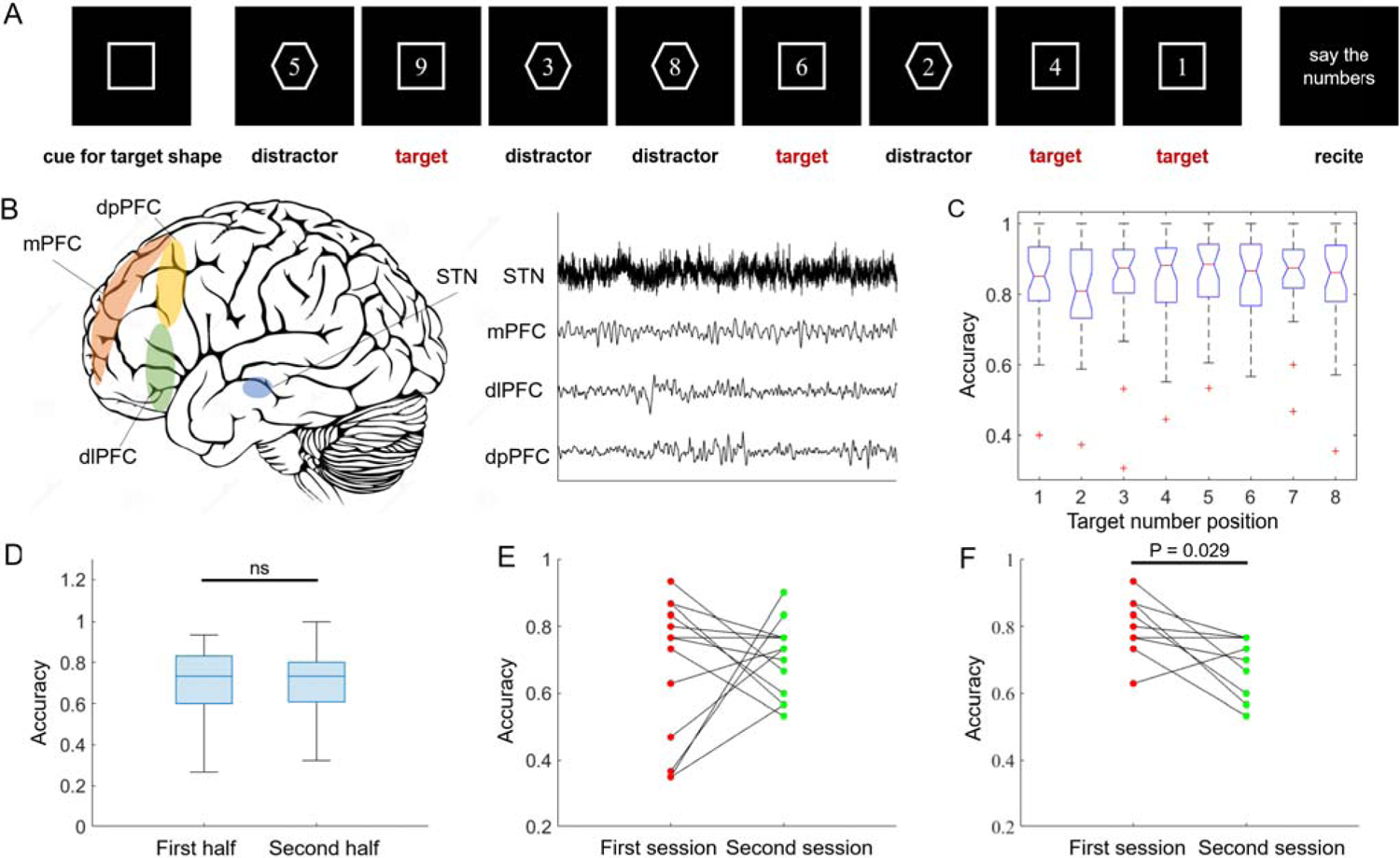
Behavior task, recording areas, and behavior results. (**A**) Non-motor decision-making task. (**B**) Left panel, ECoG strips were placed over the medial, dorsal posterior, and dorsal lateral prefrontal cortex, respectively. Macroelectrodes were placed in STN. Right panel, a sample recording from one trial is shown for electrodes placed in each region. (**C**) Accuracy for target number placed in each sequential position. (**D**) Accuracy for target numbers in first and second half o one experimental session. (**E**) Accuracy for the first and second session of one participant. (**F**) Accuracy for the first and second session of one participant, after four participants were removed.

### Time frequency analysis

We performed all analyses using custom MATLAB scripts based on fieldtrip toolbox (Oostenveld et al., 2011). The original local field potential (LFP) activity from each macroelectrode and iEEG activity from each subdural contact were bandpass filtered between 1 and 500 Hz, downsampled to 1 kHz, and notch filtered at 60 Hz. Trials that number was presented in the target shape and successfully retrieved were defined as “correct” trials. Trials that number was presented in the target shape but not retrieved at the end of block were defined as “incorrect” trials. We calculated accuracy of each session as the ratio of target trials that were successfully retrieved to all target trials. In order to get relatively balanced trials under both conditions, sessions with accuracy larger than 80% were discarded. Time frequency measures were computed as multiplication between the power spectrum of LFP signals and the power spectrum of complex Morlet wavelets, with frequency increasing from 2 to 200 Hz. We used a 1000 ms buffer on both sides of the clipped data to eliminate edge effects. We chose 500 ms preceding the presentation of each number as baseline period and calculated the percentage change in power (normalized power) by comparing the power to the mean power recorded during the baseline period. For each brain region, we average the normalized power from all electrodes within that brain region. The above procedures were repeated across trials separately for “correct” and “incorrect” trials.

### Inter trial phase coherence

Inter trial phase coherence (ITPC) is a measure of how consistent the oscillatory phase is across trials. Here, we used the Fourier spectrum achieved through the wavelet analysis to investigate phase relationships of low frequency (2-30 Hz) for each electrode across correct trials and incorrect trials. ITPC was indexed according to the following formula (Diepen et al., 2018):

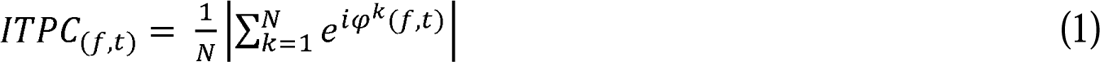

Where N is the number of trials, φ^k^ is the instantaneous phase of the signal at frequency f and time t in trial k.

### Inter-regional coherence

To quantify the time-varying coherence between two brain regions, we calculated the phase locking value (PLV) between every pair of electrodes from two brain regions at each time-frequency point. The PLV was indexed according to the following formula (Lachaux et al., 1999):

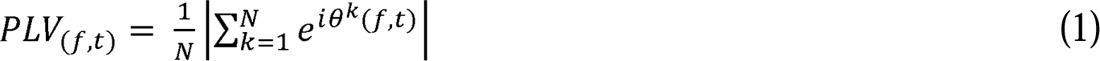

Where, θ^k^*(f, t)* is the phase difference for electrode pair (a, b), at frequency f and time t in trial k. N is the number of trials. PLV ranger from 0 to 1, with higher value if the phase differences between electrodes vary less across time.

### Granger causality

To examine the directionality of the information flow between two brain regions, we analyzed the cross-regional connectivity with spectral Granger causality. Spectral Granger causality quantifies the directional connectivity strength by testing the prediction error of the signal by introducing time series from another brain region in the frequency domain. We extracted the GC values within the low-frequency band (2-30 Hz) for electrode pairs each from one brain region across correct and incorrect trials, separately. The significance testing for the spectral GC was conducted by cluster-based non-parametric permutation methods (Tort et al., 2010).

### Phase amplitude coupling

To identify the cross-frequency coupling in each brain region, we examined modulation index (MI) between low frequencies (2-30 Hz) for phase data and BHG (80-200 Hz) for amplitude data in each electrode localized in each brain region. The calculation followed the procedure outlined by Tort and colleagues (Maris et al., 2007).

### Statistical analysis

During behavior analysis, we performed one-way ANOVA with post-hoc Tukey’s multiple comparison test for the accuracy comparison of target numbers presented in different positions. The accuracy between two sessions, two parts in one session, and PAC between theta-broadband high gamma was evaluated by two-sided paired t-test.

The comparison of theta/beta band LFP power, ITPC, and time-frequency cross-regional coherence between correct and incorrect trials were evaluated by cluster-based non-parametric permutation procedure (number of permutations was 10000, threshold of significance is 0.05).

## Results

We re-analyze electrophysiological data recorded in Parkinson’s patients during a novel non-motor decision-making task that requires participants to decide whether to encode items into working memory (Zavala et al., 2017). Eighteen patients (16 males; (mean SEM) years old) participated in the task. Local field potential (LFP) and neuronal spiking activity in the human STN and intracranial EEG (iEEG) activity from the PFC were captured. In our study, the LFP in the human STN, iEEG in dorsal posterior prefrontal cortex (dpPFC), dorsal lateral prefrontal cortex (dlPFC), and medial prefrontal cortex (mPFC) were used for analysis (Fig. 1B).

### Behavior results

During each task block, a target shape (square or octagon) was first presented to participants (Fig. 1A). Then eight single-digit numbers were sequentially presented to participants, in which four of the numbers appeared within the target shape (target trials). Four target numbers and four distract numbers were randomly interleaved. Participants were instructed only to attend to and memorize numbers presented within the target shape. At the end of each block, participants vocalized the numbers they remembered.

We first investigated if the position of the target number in the sequence affected the memorization of that number. As shown in Fig. 1C, the average accuracy of target number in all eight positions was above eighty percent, and no significant difference was found among positions (p>0.05, one-way ANOVA). Then, the accuracy of the first half and second half of all blocks in each session were compared, and no significant difference between two halves was found (p>0.05, paired t-test, Fig. 1D). This indicated that during each session, the performance of participants remained stable. Fourteen out of eighteen participants conducted the task twice, with recording captured in either left- or right-sided brain regions. We tested the accuracy between two sessions for these fourteen participants. As shown in Fig. 1E, most participants made more mistakes in the second session. But four of them performed much better in the second session. The bad performance of these four participants in the first session may be caused by maladaptive to the environment. After removal of these four participants, accuracy of remaining participants was significantly lower during the second session as compared to the first session (paired t test: t(9) = 2.5866, p=0.029; Fig. 1F).

### Attention-related neural activity in STN

Studies based on fMRI functional connectivity analysis, encephalographic (EEG) event-related potential (ERP) and repetitive TMS (rTMS) have identified dorsal lateral prefrontal cortex may serve as source of top-down modulation underlying visual attention (Zanto et al., 2010; Zhang et al., 2008; Zanto et al., 2011). Human EEG and MEG studies showed increases in theta (3-7 Hz) activity in dorsal and ventral frontal regions, and anterior cingulate cortex (ACC) linked to attentional amplification ((McDermott et al., 2017; Szczepanski et al., 2014). To test this, we analyzed the local field potential (LFP) recorded in dpPFC, dlPFC, mPFC, and STN while patients performed a novel non-motor decision-making task (Fig. 2A). The average power for each electrode was computed separately across all correct and incorrect trials during each session. In line with previous studies, theta power in dlPFC showed significant increases during the correct trials than incorrect trials, suggesting increased theta power related to sustained attention (Fig. 2B). Other regions exhibited tendency to initially increase and then decrease in theta power under both conditions, but the differences between conditions were not significant (Fig. 2B). Cross all sessions, beta power in dpPFC and dlPFC exhibited decreases following the presentation of stimulus under both conditions (Fig. 2C). The beta power in STN remain stable during the correct trials, and first increased then decreased during the incorrect trials (Fig. 2C).

**Fig. 2.**
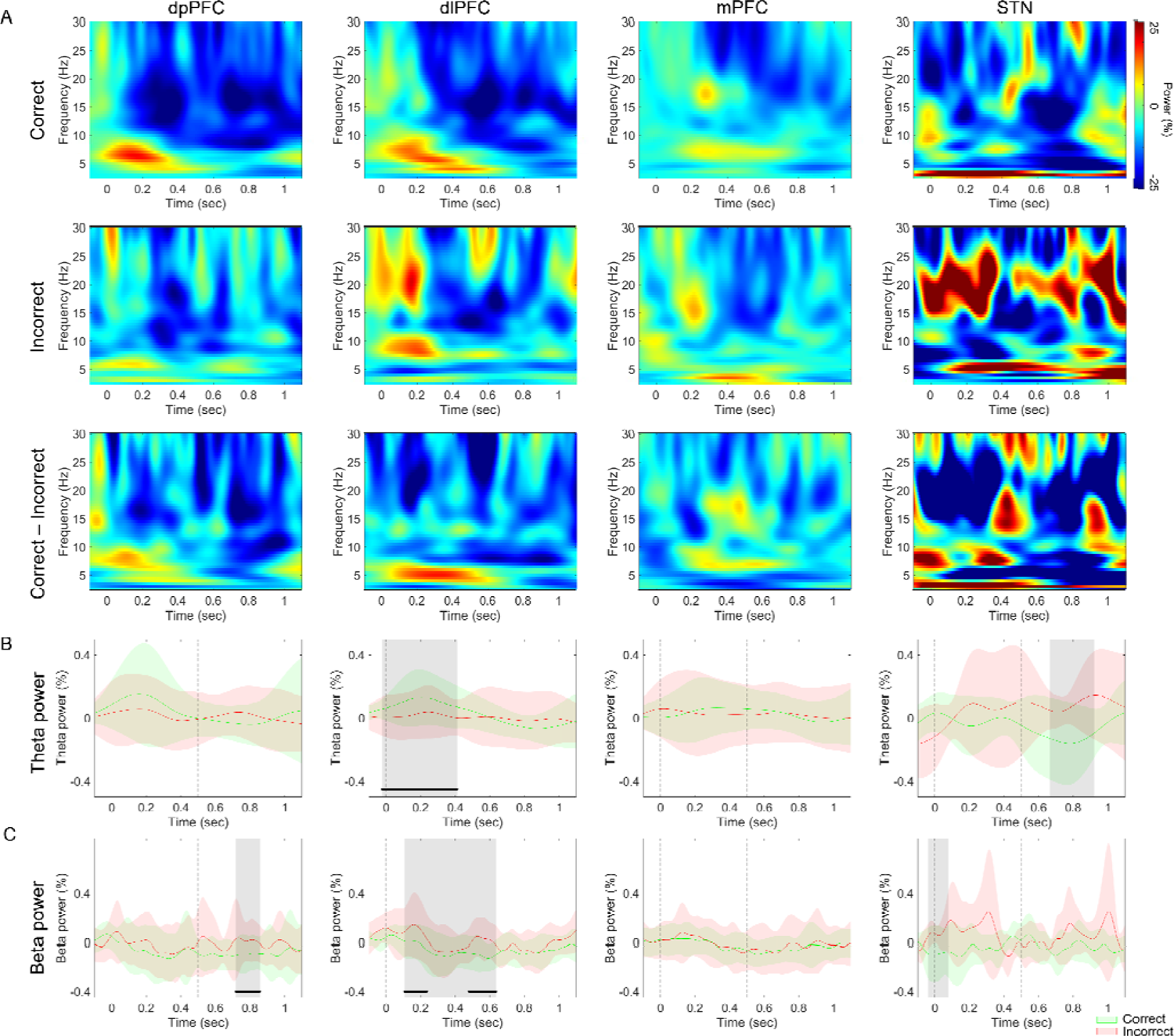
power spectrum between correct and incorrect trials. (**A**), Normalized oscillatory power averaged across all dpPFC electrodes (left), all dlPFC electrodes (left-middle), all mPFC electrodes (middle-right), and all STN electrodes (right). In each brain region, the power was averaged across correct trials (top), incorrect trials (middle), and correct minus incorrect (bottom). Stimulus onset time is denoted by time point zero on the X-axis. (**B**), Normalized oscillatory power averaged across electrodes located in each brain region, across theta band. Time points exhibiting a significant difference between each condition (p < 0.05, cluster-based permutation test) are denoted by black horizontal bar. Time points exhibiting a tendency to show differences between each condition (p < 0.1, cluster-based permutation test) are denoted by gray shaded box. Stimulus onset time is denoted by time point zero on the X-axis. (**C**), Same as (B), but averaged across the theta band.

Some researchers investigate the effect of attention on inter-trial phase coherence at pre-stimulus low-frequency oscillations in area MT of monkey. Their results reveal that phase coherence in theta band increases when attention is deployed towards the receptive field of the recorded neuron (Zareian et al., 2020). In our study, we compared inter-trial phase coherence (ITPC) between the correct and incorrect trials in frontal regions and STN (Fig. 3). The ITPC during incorrect trials (Fig. 3A, left panel) was significantly higher than correct trials (Fig. 3A, middle panel), indicated that sustained attention related to decreased ITPC in these regions. Cluster-based statistics confirmed statistically significant clusters, comprising delta, theta, alpha, and beta band in three prefrontal regions, and theta, alpha, and beta band in STN (Fig. 3A, right panel). The clusters in three prefrontal regions were last the whole trial duration, and the clusters in STN were spread in the late part of trial duration.

**Fig. 3.**
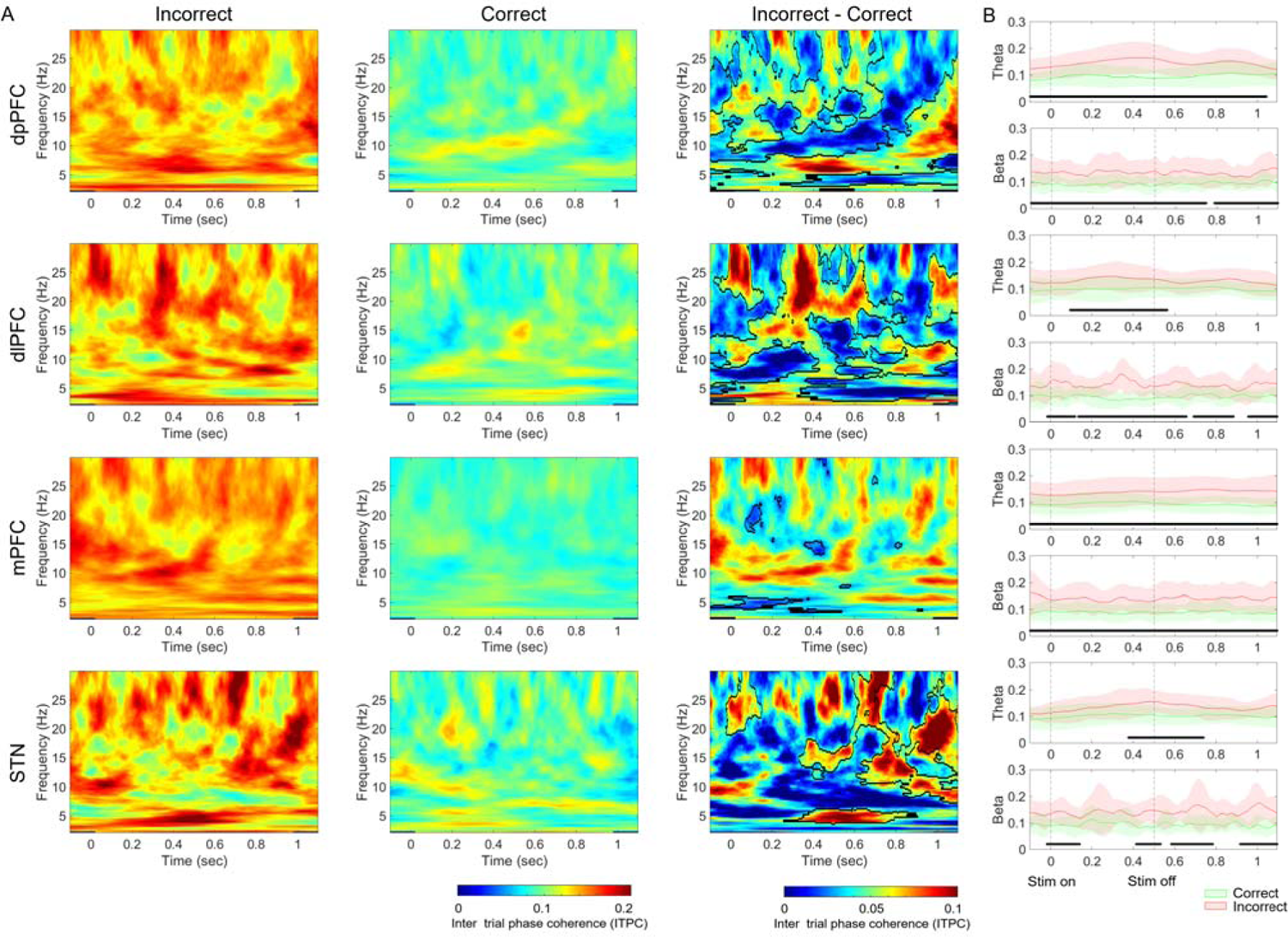
Inter trial phase coherence (ITPC) between correct and incorrect trials. (**A**), ITPC averaged across all dpPFC electrodes (top), all dlPFC electrodes (top-middle), all mPFC electrodes (middle-bottom), all STN electrodes (bottom). In each brain region, the ITPC was averaged across incorrect trials (left), correct trials (middle), and incorrect minus correct trials (right). Mask in the right panel denotes time-frequency regions exhibiting significant differences between conditions (p < 0.05, cluster-based permutation test). Stimulus onset time is denoted by time point zero on the X-axis. (**B**), Same as (A), but averaged across the beta and theta band. Time points exhibiting a significant difference between each condition (p < 0.05, cluster-based permutation test) are denoted by black horizontal bar.

Given the important role of theta and broadband high gamma (BHG) oscillations in attentional processing (Baird et al., 2014; Foster et al., 2013; Helfrich et al., 2018)). We assessed whether any similar relationship existed among the different frequency bands (delta, theta, alpha, beta, high gamma) during attentional fluctuations in human prefrontal regions and STN. We examined PAC between a wide range of frequencies for phase data (2-30 Hz), and BHG (a proxy for local spiking activity or cortical excitability (Ray et al., 2011; Watson et al., 2017)) for amplitude data (80-200 Hz) in all of the electrodes spread across four regions (Fig. 4A). We found significant coupling between the phase of the delta/theta signal (2-8 Hz) and BHG amplitude (80-200 Hz) during incorrect trials compared to correct trials (Fig. 4B, C). We also observed alpha phase-BHG amplitude coupling in electrodes over dpPFC (Fig. 4A, right panel, top). Our results indicated that decreased theta-BHG coupling related to sustain attention, complied with previous nonhuman primates studies (Spyropoulos et al., 2018; Esghaei et al., 2015).

**Fig. 4.**
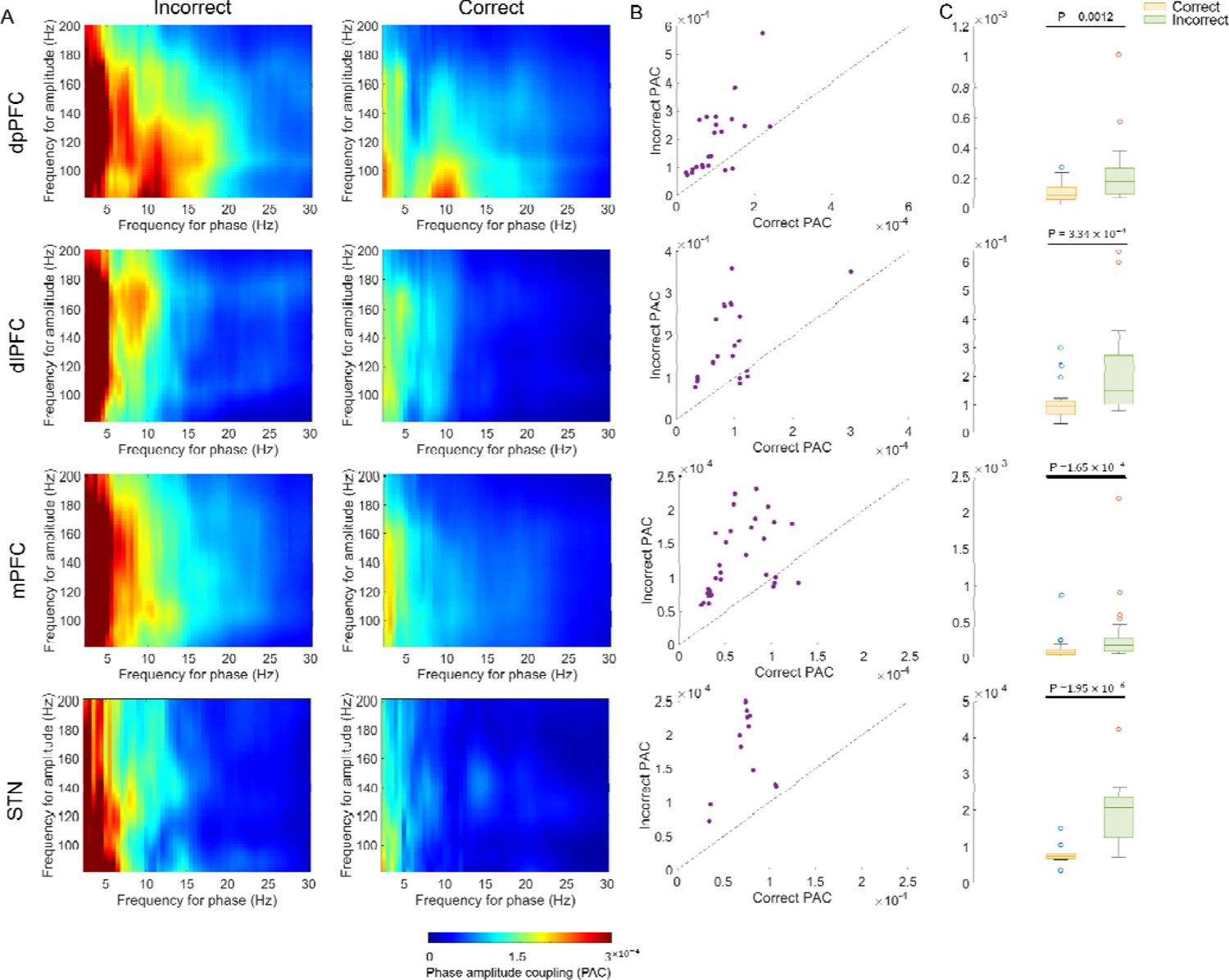
Phase amplitude coupling (PAC) between theta and BHG. (**A**) PAC averaged across all dpPFC electrodes (top), all dlPFC electrodes (top-middle), all mPFC electrode pairs (middle-bottom), and all STN electrodes (bottom). In each brain region, the PAC was averaged across incorrect trials (left) and correct trials (right). (**B**) Individual electrode’s PAC values for correct and incorrect trials. Each circle represents data from a single electrode with PAC calculated separately for correct trials (X-axis) and incorrect trials (Y-axis). (**C**) Bar plot showing group average of PAC values over theta phase frequencies (4-8 Hz) and BHG amplitude frequencies (80-200 Hz) for correct and incorrect trials, across electrodes in each brain region. Error bars represent standard deviation.

### Attention-related STN-PFC circuit connectivity

We next examined the coherence between areas. We found the coherence during incorrect trials was higher than during correct trials among all STN-PFC area pairs (Fig. 5, left and middle panels). The coherence between dpPFC and STN was significantly different in delta, theta, alpha, and beta bands across the whole trial duration (p < 0.05, two-sided permutation test; Fig. 5, right panel). The coherence between dlPFC and STN was significantly different in theta, alpha, and beta band, beginning at 100ms after stimulus presentation and last several hundred milliseconds after stimulus disappear (p < 0.05, two-sided permutation test; Fig. 5, right panel). The coherence between mPFC and STN was significantly different in theta, alpha, and beta bands, during the whole trial. The coherence between prefrontal regions was significantly different in alpha and beta bands, during the whole trial duration (p < 0.05, two-sided permutation test; Fig. 5, right panel). The coherence between pairs of PFC sub-regions was shown in supplement figure 1. The same tendency was shown between PFC sub-regions that, coherence during incorrect trials was higher than during correct trials.

**Fig. 5.**
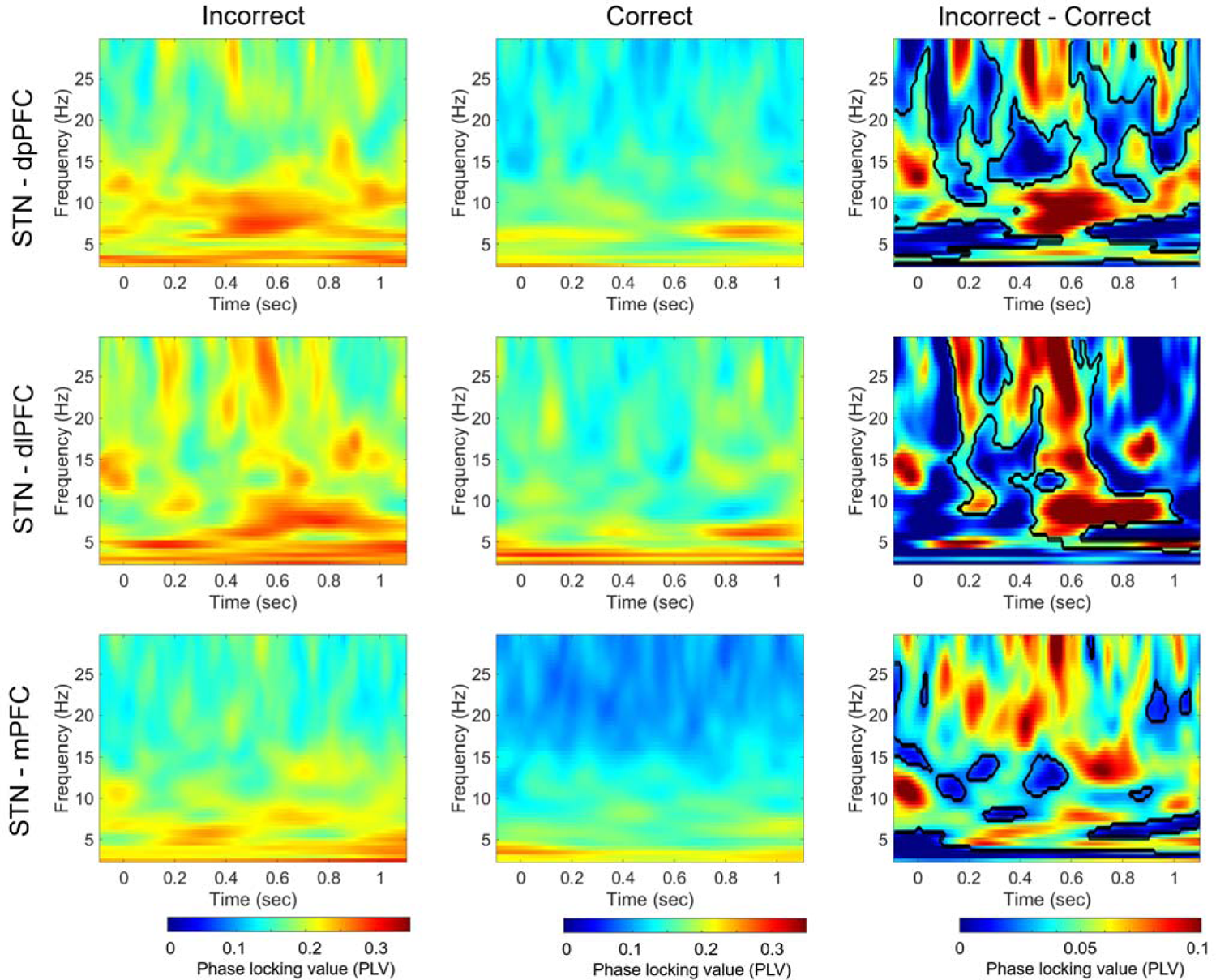
Phase locking value (PLV) between STN and prefrontal areas. PLV averaged across all STN-dpPFC electrode pairs (top panel), all STN-dlPFC electrode pairs (middle panel), all STN-mPFC electrode pairs (bottom panel). In each brain region, the PLV was averaged across incorrect trials (left), correct trials (middle), and incorrect minus correct trials (right). Mask in the right panel denotes time-frequency regions exhibiting significant differences between conditions (p < 0.05, cluster-based permutation test). Stimulus onset time is denoted by time point zero on the X-axis.

As PAC also serves as an important mechanism for inter-regional information transfer, we computed the cross-regional PAC in both phase-amplitude combinations between all four regions, for both correct trials and incorrect trials (Fig. 6A). The inter-regional PAC was quantified as the coupling between the low-frequency phases (2-30 Hz) from the electrode in one region and the high-frequency amplitude (80-200 Hz) from the electrode in another region. The results showed that the delta/theta-BHG coupling was different between correct trials and incorrect trials for all region pairs (Fig. 6A, D, G). To further compare the coupling strength between different trials in all regional combinations, the mean of cross-regional PAC of the delta/theta-BHG bands was extracted. With post hoc paired t-test evaluation, we found that the coupling strength during the incorrect trials was significantly greater than during the correct trials (Fig. 6B, C, E, F, H, I). PAC across prefrontal sub-regions were shown in supplement figure 2.

**Fig. 6.**
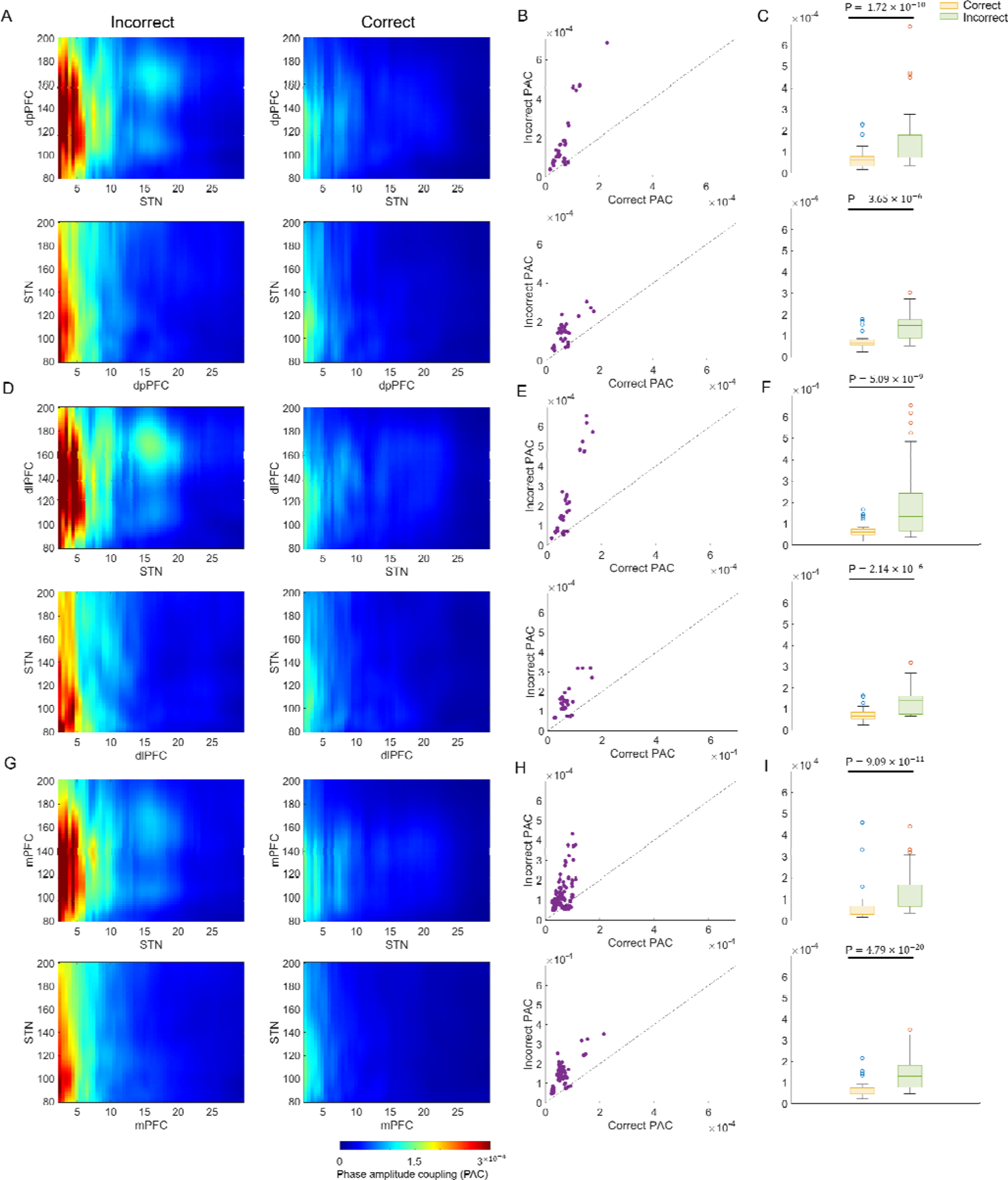
Cross-regional phase amplitude coupling between STN and prefrontal areas. (**A**) PAC for low-frequency phase from STN and BHG frequency amplitude from dpPFC averaged across all electrode pairs (top), for low-frequency phase from dpPFC and BHG frequency amplitude from STN averaged across all electrode pairs (bottom). For each coupling pair, the PAC was averaged across incorrect trials (left) and correct trials (right). (**B**) Individual electrode pair’s PAC values for correct and incorrect trials. Each circle represents data from a single electrode pair with PAC calculated separately for correct trials (X-axis) and incorrect trials (Y-axis). (**C**) Bar plot showing group average of PAC values over theta phase frequencies (4-8 Hz) and BHG amplitude frequencies (80-200 Hz) for correct and incorrect trials, across electrode pairs across brain regions. Error bars represent standard deviation. (**D**) same as (A), but between STN and dlPFC. (**E**) same as (B), but between STN and dlPFC. (**F**) same as (C), but between STN and dlPFC. (**G**) same as (A), but between STN and mPFC. (**H**) same as (B), but between STN and mPFC. (**I**) same as (C), but between STN and mPFC.

Our results showed that sustained attention was associated with decreased connectivity between STN and prefrontal regions, and communication across brain areas was achieved through different oscillation bands. The theta band played an important role in communication between frontal regions and STN.

### Task related information flow across STN-PFC circuit

In order to uncover the direction of functional interactions within each area pair, we adopted Granger causality analysis. We found significant STN leading information flow to dlPFC in theta and alpha band (P<0.05, paired t-test; Fig. 7A), to dpPFC in theta band (p<0.05, paired t-test; Fig. 7B), and to mPFC in high theta, alpha and beta band (P<0.05, paired t-test; Fig. 7C), across all trials (correct and incorrect trials) during the task. The information flow from dpPFC to STN in high beta band (p<0.05, paired t-test; Fig. 7B), from mPFC to STN in delta band (p<0.05, paired t-test, Fig. 7C). We observed significant information from dlPFC to dpPFC in alpha band, from mPFC to dpPFC in delta and theta band, and from dlPFC to mPFC in alpha band (p<0.05, paired t-test; supplement figure 3). No significant difference was found between correct and incorrect trials, indicating that the direction of information flow was modulated by engagement of cognitive task rather than attentional fluctuation.

**Fig. 7.**
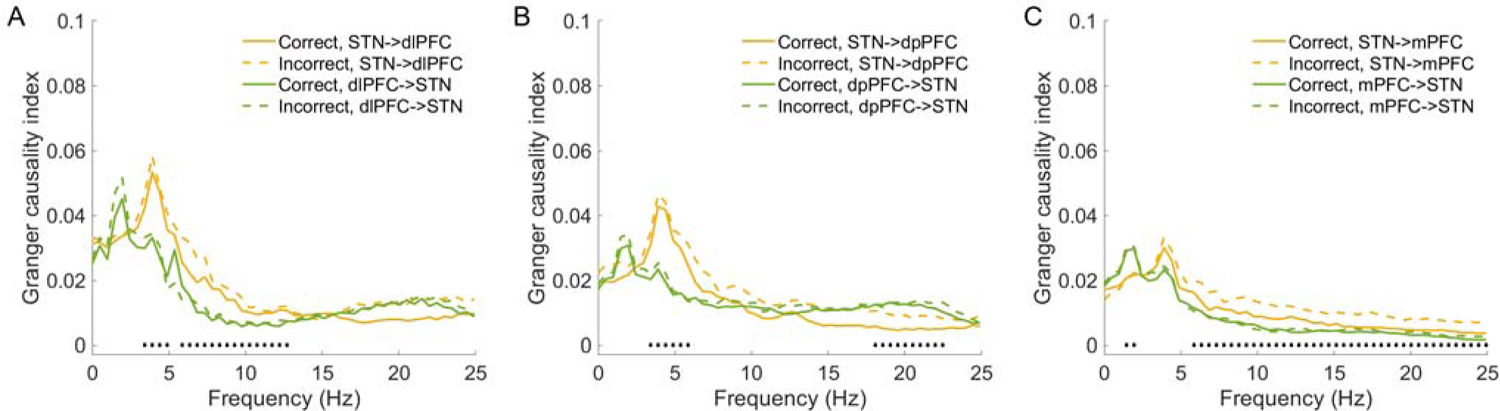
Information flow between STN and prefrontal areas. (**A**), Granger causality index across all STN-dlPFC electrode pairs. (**B**), Granger causality index across all STN-dpPFC electrode pairs. (**C**), Granger causality index across all STN-mPFC electrode pairs. Black horizontal dots denote the frequencies that show significant differences between two opposite information transfer directions (p < 0.05, paired t-test).

## Discussion

Researchers used LFP power recorded from area MT of macaque to decode the context the animal was attending to. Both delta/theta (<9 Hz) and broadband gamma (31-120 Hz) activities contain sufficient information to decode (Esghaei et al., 2014). By examining LFP data from large-scale ECoG recordings, researchers revealed that attentional allocation and behavioral performance fluctuate as a function of the theta phase in widespread regions of the frontoparietal attention network. They hypothesized that low-frequency oscillations are a direct result of periodic fluctuations and cortical high-frequency band (HFB) is modulated by the low-frequency activity (Helfrich et al., 2018). In our study, we found STN activity covaried with attentional fluctuation, too. ITPC in the STN during correct trials is significantly lower than during incorrect trials across theta, alpha, and beta bands, in line with previous brain-computer interface (BCI) paradigms in EEG (Zareian et al., 2020). The theta-BHG coupling in prefrontal regions and STN was also covaried with attentional fluctuation. The theta-BHG PAC was significantly higher during the incorrect trials than during the correct trials. As BHG activities are sufficient proxy for local spiking activities, our results were in line with previous study performed on macaque monkey (Spyropoulos et al., 2018; Esghaei et al., 2015), in which attentional de-synchronization was mediated by de-coupling between local spiking activities and low-frequency oscillations (Fries et al., 2001; Buffalo et al., 2011; Esghaei et al., 2018).

Electrophysiological studies have shown that low-frequency interactions between PFC and STN support cognitive control, where fronto-STN circuits pause behavior and cognition when stop signals or conflict signals occur (Frank et al., 2007; Aron et al., 2016; Zeng et al., 2021). Theta activity of the PFC was shown to drive the activity of the STN, which adjusted the threshold of evidence in a dot motion discrimination task (Zavala et al., 2014). The STN DBS disrupted the correlation between PFC theta power and behavior responses (Cavanagh et al., 2011). The information flow seemed to be bidirectional, as a human intracranial study showed that STN stimulation evoked short-latency lateral PFC and motor cortical responses (∼12 ms) by using electrical stimulation tract-tracing (ESTT) (Kelley et al., 2017). The same study also revealed 4-Hz Granger causal connectivity from STN to midfrontal EEG during conflict tasks, and 4-Hz STN DBS improved cognitive performance (Kelley et al., 2017).

In our study, we found STN activity covaried with attentional fluctuation and STN-PFC circuits support this process. The coherence between STN and frontal regions was significantly higher during the incorrect trials than correct trials. Our cross-region PAC analysis also showed that the coupling between theta phase from one region and high gamma amplitude from another region was significantly higher during the incorrect trials than correct trials. As low-frequency coherence between regions was significantly higher during incorrect trials, we believe the cross-regional PAC coupling was mediated by the coherent cross-regional low-frequency. GC analysis showed significant theta band connectivity from STN to prefrontal regions, we believed that theta activity in STN drove the activity in frontal regions during cognitive tasks.

Our study was performed on prefrontal regions and STN. All of these areas are related to higher-order cognitive control. As attention is a meta modulation process between sensory areas and high-order cognitive control areas (Buffalo et al., 2011; Fries et al., 2008; Chalk et al., 2010; Mitchell et al., 2009; Cohen et al., 2009; Averbeck et al., 2006), further research on the relationship between sensory areas, prefrontal regions and STN could be very insightful.

## Acknowledgments

This work was supported by the Science Center of National Natural Science Foundation of China “Cognitive Computing” (No. 62088102), Major Project of New Generation artificial intelligence of Ministry of Science and Technology, (No. 2020AAA0105500). We thank Jiamin Wu and Qionghai Dai for the supervision of the whole project and help in project design and manuscript preparation. We thank Sen Wan and Yanwang Zhai for very fruitful discussions.

---

**Supplement Fig. 1.**
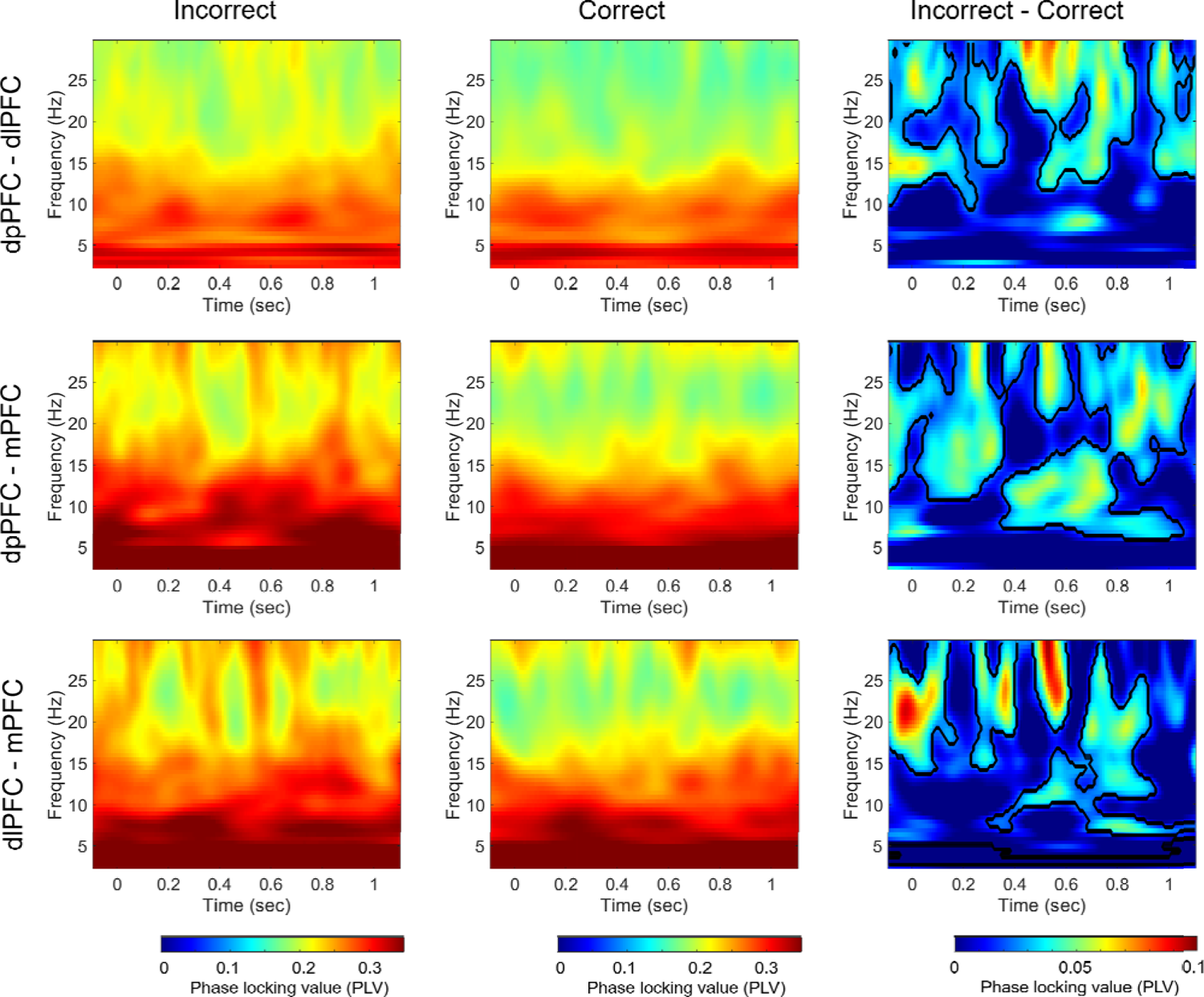
Phase locking value (PLV) across prefrontal areas. PLV averaged across all dpPFC-dlPFC electrode pairs (top panel), all dpPFC-mPFC electrode pairs (middle panel), all dlPFC-mPFC electrode pairs (bottom panel). In each brain region, the PLV was averaged across incorrect trials (left), correct trials (middle), and incorrect minus correct trials (right). Mask in the right panel denotes time-frequency regions exhibiting significant differences between conditions (p < 0.05, cluster-based permutation test). Stimulus onset time is denoted by time point zero on the X-axis.

**Supplement Fig. 2.**
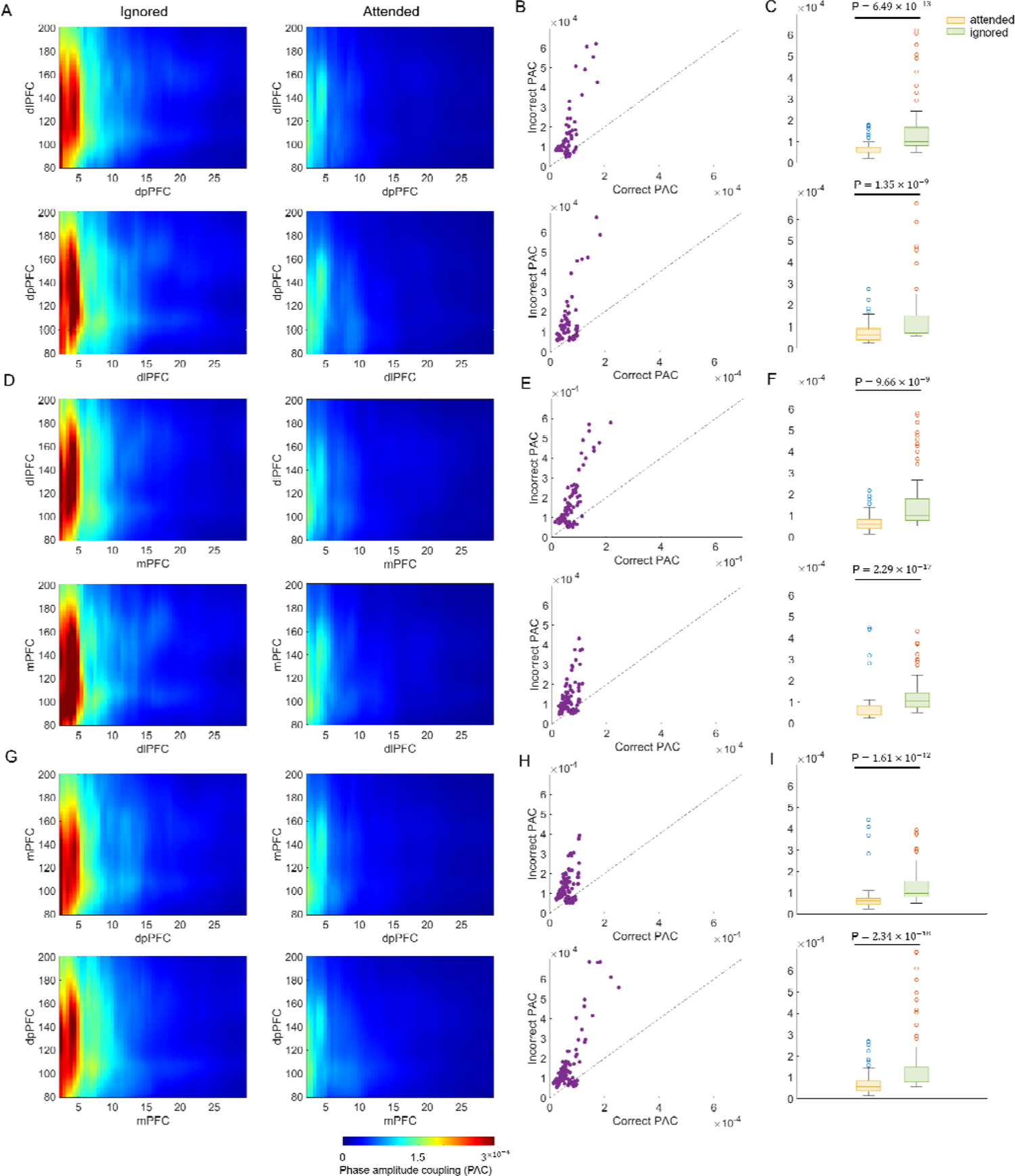
Cross-regional phase amplitude coupling across prefrontal areas. (A) PAC for low-frequency from dpPFC and BHG frequency amplitude from dlPFC averaged across all electrode pairs (top), for low-frequency hase hase from dlPFC and BHG frequency amplitude from dpPFC averaged across all electrode pairs (bottom). For each coupling pair, the PAC was averaged across incorrect trials (left) and correct trials (right). (B) Individual electrode pair’s PAC values for correct and incorrect trials. Each circle represents data from a single electrode pair with PAC calculated separately for correct trials (X-axis) and incorrect trials (Y-axis). (C) Bar plot showing group average of PAC values over theta phase frequencies (4-8 Hz) and BHG amplitude frequencies (80-200 Hz) for correct and incorrect trials, across electrode pairs across brain regions. Error bars represent standard deviation. (D) same as (A), but between mPFC and dlPFC. (E) same as (B), but between mPFC and dlPFC. (F) same as (C), but between mPFC and dlPFC. (G) same as (A), but between dpPFC and mPFC. (H) same as (B), but between dpPFC and mPFC. (I) same as (C), but between dpPFC and mPFC.

**Supplement Fig. 3.**
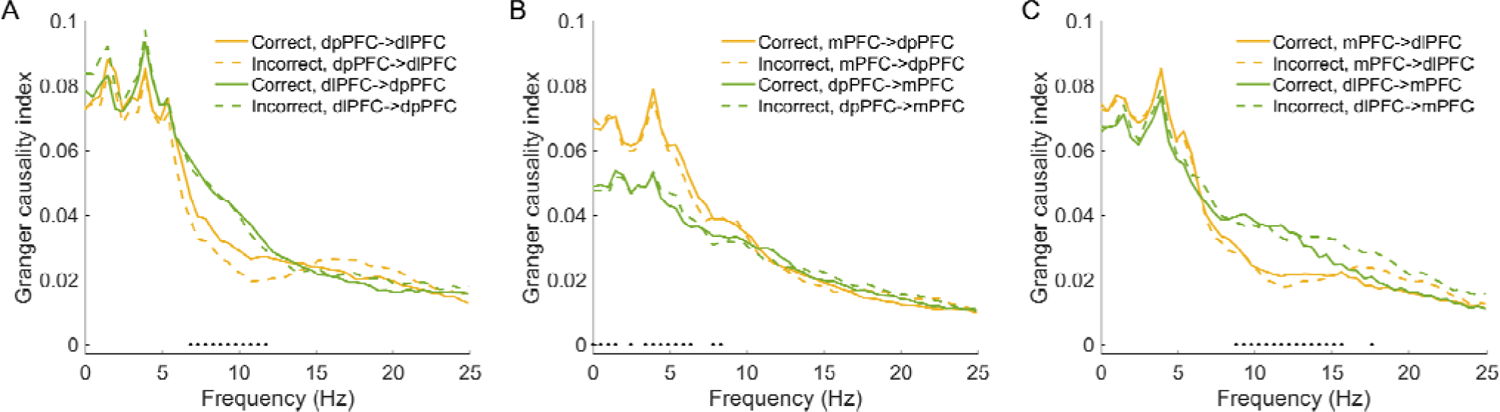
Information flow across prefrontal areas. (A), Granger causality index across all dpPFC-dlPFC electrode pairs. (B), Granger causality index across all mPFC-dpPFC electrode pairs. (C), Granger causality index across all dlPFC-mPFC electrode pairs. Black horizontal dots denote the frequencies that show significant differences between two opposite information transfer directions (p < 0.05, paired t-test).

